# Gaze dynamics of feature-based distractor inhibition under the prior-knowledge and expectation

**DOI:** 10.1101/2020.07.19.210781

**Authors:** Wen Wen, Yangming Zhang, Sheng Li

**Affiliations:** School of Psychological and Cognitive Sciences and Beijing Key Laboratory of Behavior and Mental Health, Peking University, Beijing, China; PKU-IDG/McGovern Institute for Brain Research, Peking University, Beijing, China; Key Laboratory of Machine Perception (Ministry of Education), Peking University, Beijing, China

**Keywords:** feature-based attention, suppression, expectation, decoding

## Abstract

Prior information about distractor facilitates selective attention to task-relevant items and helps the optimization of oculomotor planning. Particularly, feature-based attentional inhibition could be benefited from the pre-knowledge of critical features of the distractors. In the present study, we capitalized on gaze-position decoding to examine the dynamics of attentional deployment in a feature-based attentional task that involved two groups of dots (target/distractor dots) moving toward different directions. Specifically, this measurement revealed how pre-knowledge of the target’s or distractor’s direction modulated real-time feature-based attentional bias. In Experiment 1, participants were provided with target cues indicating the moving direction of target dots. The results showed that participants were biased towards the cued direction and tracked the target dots throughout the task period. In Experiment 2 and Experiment 3, participants were provided with cues that informed the moving direction of distractor dots. The results showed that participants would continuously monitor the distractor’s direction when the distractor cue varied on a trial-by-trial basis (Experiment 2). However, when the to-be-ignored distractor direction remained constant (Experiment 3), participants would strategically bias their attention to the distractor’s direction before the cue onset and reduce the cost of re-deployment of attention between trials. These results suggest that monitoring the distractor’s feature is a prerequisite for feature-based attentional inhibition and this process is facilitated by the predictability of the distractor’s feature.

## Introduction

To process the rich and fast-changing visual input, human needs to efficiently assign the limited attentional resource to task-relevant stimuli. Target cues have been proven to be beneficial in promoting target selection (Posner, 1980; Vickery, King, & Jiang, 2005; Wolfe, 1994). However, whether prior knowledge of distractors (i.e., distractor cues) can help us to avoid unnecessary attention allocation to task-irrelevant information and facilitate performance is still an issue under debate. Divergent findings have been reported regarding whether distractor cues can accelerate search efficiency (Arita, Carlisle, & Woodman, 2012; Beck & Hollingworth, 2015; Becker, Hemsteger, & Peltier, 2015; Olivers, 2009; Soto, Heinke, Humphreys, & Blanco, 2005; Geoffrey F Woodman & Luck, 2007). Among the studies that showed behavioral promotions, the emergence of suppression benefits seemed to be dependent on multiple factors (Conci, Deichsel, Müller, & Töllner, 2019; Han & Kim, 2009; Stilwell & Vecera, 2019; Tanda & Kawahara, 2019; Töllner, Conci, & Müller, 2015), especially the constancy of distractor cues across trials so as to form expectations (Cunningham & Egeth, 2016; Gaspelin, Leonard, & Luck, 2015; Gaspelin & Luck, 2018a; Vatterott & Vecera, 2012; Wen, Hou, & Li, 2018, for reviews, see Gaspelin & Luck, 2018b, 2019; Noonan, Crittenden, Jensen, & Stokes, 2018; van Moorselaar & Slagter, 2020).

Despite the increasing number of studies on whether the distractors can be efficiently inhibited, the question of how do we filter out them remains untangled. Researchers have proposed two possible mechanisms: proactive vs. reactive suppression (Geng, 2014) Supporting evidence has been observed for both mechanisms. Some studies found that rejection templates would be created based on the foreknowledge of distractors and proactively guide our attention away from matched stimuli (Woodman, Carlisle, & Reinhart, 2013; Woodman & Luck, 2007). For example, a salient color-singleton distractor can be proactively inhibited and letters inside of the singleton distractor were less likely to be reported (Gaspelin et al., 2015). Moreover, an eye-tracking study showed that fewer gazes were directed to the color-singleton distractor than other search items (an oculomotor suppression effect) and this occurred even when the eye movements were initiated relatively quick, indicating that salient items can be proactively suppressed (Gaspelin, Leonard, & Luck, 2017). Event-related-potential (ERP) evidence demonstrated that salient distractors evoked the Pd component to intervene in the automatic capture (Gaspar & McDonald, 2014; Sawaki & Luck, 2011). Hence, the *signal suppression hypothesis* suggested that the suppression of the salient item occurs before the initial capture (Gaspelin & Luck, 2018b; Sawaki & Luck, 2011).

On the contrary, some researchers contended that distractor suppression is a reactive process and suppression cannot occur unless the distractor has been attended to. For instance, Moher and Egeth (2012) provided distractor cues before the search array and participants were slower in the negative-cue condition than in the neutral condition. By manipulating the stimulus onset asynchrony (SOA), they found the trend of avoidance in long SOA trials. Hence, they proposed the *search and destroy* process where attention is biased toward matched distractors and then rejected given sufficient time. This reactive inhibition hypothesis echoed the literature that proposed the mechanism of rapid disengagement (Awh, Belopolsky, & Theeuwes, 2012; Theeuwes, 2010). Liesefeld and colleagues (2017) further examined the temporal dynamics of Pd and N2pc evoked by the distractor and target to demonstrate the disengagement from the misallocated attention to the salient distractor and re-direction of attention to the target.

Critically, the majority of the existing studies changed the distractor feature from trial to trial, which hampered the establishment of stable inhibition templates through learning. Recent findings seemed to reach the consensus that the ability to inhibit distractors can be learned (Gaspelin & Luck, 2019; Geng, Won, & Carlisle, 2019) when the distractor feature remained constant or when the distractor locations were statistically manipulated (Wang & Theeuwes, 2018). Under such arrangements, the initial capture would disappear and the rejection benefit would emerge gradually along with blocks (Cunningham & Egeth, 2016; Stilwell & Vecera, 2019; Vatterott & Vecera, 2012).

Most of the above-mentioned studies used a visual search paradigm where items were spatially separated. Although the participants were informed of the distractor feature using the distractor cue, they would transform the feature cue into a spatial filter so as to suppress the matched items. One dominant discrepancy between feature-based attention and spatial attention is that feature-based attention does not rely on specific locations. Thus, common oculomotor events, such as fixation, dwell time and first saccades, which are location-based, are not effective indicators of attentional allocation for features that are spatially inseparable. To unequivocally reveal how distractor inhibition evolves over the course of a trial, we need a finer-scale measurement that can provide continuous information about attentional deployment at a millisecond level.

In the present study, we aimed to investigate the role of prior information in feature-based attentional inhibition. To avoid the confounding factors that are related to spatial attention, we used moving dots stimuli that were intermingled and spatially inseparable. Participants were asked to fixate at the center of the stimuli and their eye positions were recorded during the task. In the task, participants were instructed to detect speed change in one of the two moving directions. Previous literature has suggested that small fixational gaze shifts could reflect covert attention to external stimuli (Engbert & Kliegl, 2003; Hafed & Clark, 2002) or internal attentional selection inside working memory (van Ede, Chekroud, & Nobre, 2019). These attributes make small gaze shift an ideal measurement of attention for the stimuli and task used in the present study. Specifically, we examined the gaze positions using a decoding approach to inform the temporal dynamics of attentional deployment when participants received target cues (Experiment 1), distractor cues (Experiment 2), and repeated distractor cues (Experiment 3).

## Experiment 1

### Participants

Eighteen participants were recruited in Experiment 1 and were paid for their participation. All participants (including those in Experiments 2 and 3) had a normal or corrected-to-normal vision and had no history of psychiatric or neurological disorders. They (including participants in Experiments 2 and 3) provided written informed consent prior to the experiment.

One participant’s eye data was missing due to technical failure. One participant showed extremely bad performance (detection sensitivity was more than 1.5 SD below the mean and response time was 3 SD beyond the mean). These two participants were excluded from the analysis. The final sample consisted of sixteen participants (mean age = 21.3, range from 19 to 26, 5 males, all right-handed). The study was approved by the local ethics committee at Peking University.

### Method

The stimuli were presented using MATLAB (Mathworks, Natick, MA, USA) with Psychtoolbox3 extensions (Brainard, 1997) and were displayed on a LED monitor (refresh rate: 100Hz, resolution: 2560×1440) with a viewing distance about 80 cm. As shown in Fig. 1A, each trial began with a cue indicating the moving direction of the target dots. The cue was presented for 0.3s. During the delay period, a set of dots (N = 260, 0.12° × 0.12°, speed = 3 deg/s, lifetime = 0.1s) appeared in an aperture (outer circle diameter = 20°, inner circle diameter = 8°) centered at the fixation cross (0.4°) and moved randomly for 0.6s. Once the grey fixation cross changed into green, the dots started to move coherently (coherence level = 0.8) toward two different directions. Dots in the target direction (target dots, N = 130) would move toward the cued direction while the distractor dots would move toward another direction. The directions were randomly and independently selected from eight direction bins which covered the whole circle. It was possible that the moving directions of target dots and distractor dots belonged to the same direction bin but the minimal difference must be above 15°. In half of the trials, the speed of the target dots would increase (two times upon the speed) and the increment lasted for 0.3s. The speed-up might start at any time from 0.3~1.3s after the onset of the central green fixation. Participants were asked to detect the speed-up and respond whenever it happened. The moving dots disappeared immediately after the response. For the no-change trials, coherent moving would be displayed for 1.5s. Participants completed 16 blocks of 80 trials and they received feedback about their performance at the end of each block.

**Figure 1.**
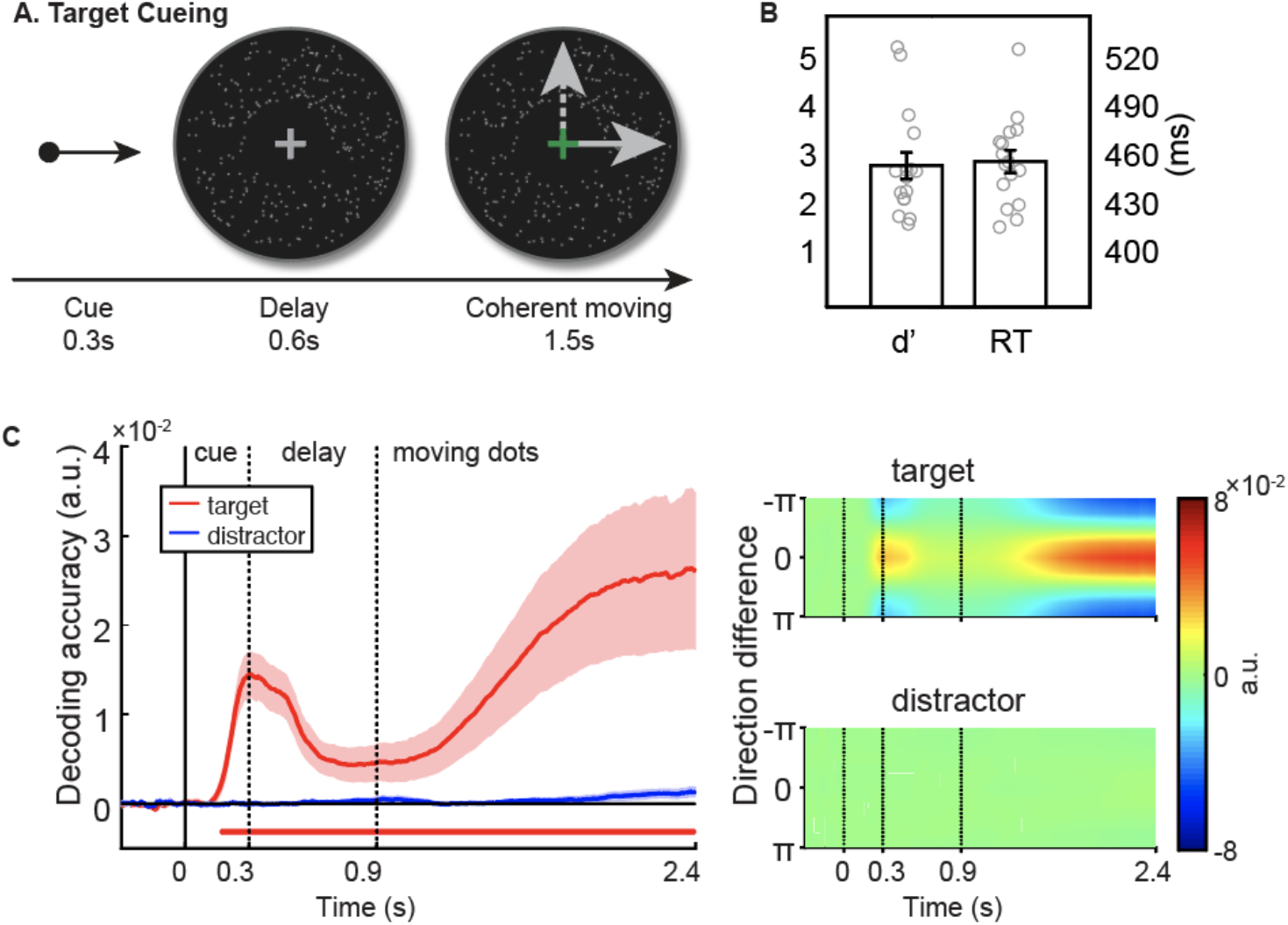
Task and results of Experiment 1. (A)Task. The cue indicated the moving direction of the target dots. All dots moved randomly during the delay. Participants need to detect the speed change of target dots during the coherent moving period. The solid/dashed arrow represents the moving direction of the target/distractor dots which were plotted for illustrations. (B) Behavioral performance. Error bars represented standard errors of the mean. Each circle dot represented one participant. (C) Direction decoding using gaze positions. The left panel was the time series of decoding accuracy. Shaded errorbar was the standard deviation of mean derived from 5,000 bootstrapped samples. Solid lines below the x-axis are the cluster-permuted significant time intervals. The right panel was the tuning curve of the averaged eight directions for the target and distractor direction at each timepoint. The tuning curve was the mean-centered and sign-reversed Mahalanobis distance between patterns to all moving directions.

Eyelink 1000 Plus (SR Research) was used to record the binocular movements with a sample rate of 500Hz. At the beginning of each block, the eye tracker was calibrated using a five-point calibration procedure. Participants were asked to fixate at the central fixation cross and avoid tracking the moving dots during the whole experiment. Their heads were stabilized using a chin-rest.

### Data analysis

#### Behavior

To compute the detection sensitivity (d’), we took trials in which responses were later than the speed change onset as hit. Standard corrections were performed to deal with hit rate of 1 or false alarm rate of 0 (Macmillan & Kaplan, 1985). Reaction time was measured as the time between the speed change onset and the response. Mean RT was obtained from hit trials.

#### Decoding

We carried out the decoding analysis using the correct-rejection trials. Eye blinks were spline-interpolated (van Ede et al., 2019). Given that the gaze position of the left and right eye should be highly correlated, we averaged the binocular data and obtained the two-dimensional data representing the raw horizontal and vertical gaze positions. We epoched the continuous data −0.3~2.4s time-locked to the cue onset and baseline-corrected to the mean of −0.3~0s before cue onset. Finally, we smoothed the time-series data using a Gaussian kernel (standard deviation = 4ms). Decoding was performed on each time point based on the Mahalanobis distance in a leave-one-out manner. Trials were randomly partitioned into 7 training-folds and 1 test-fold. To create an unbiased training set, the number of trials from each direction bin was equalized by subsampling. The subsampled trials of each direction bin were averaged and convolved with a half cosine basis set raised to the 7th power. The covariance matrix was estimated based on the convolved training set. The Mahalanobis distances were computed between each trial of the test set and the averaged basis-weighted training set. After obtaining the Mahalanobis distances, we mean-centered them across the eight directions. The distances to the eight directions were regarded as the ‘tuning curve’. By computing the cosine vector mean of the tuning curve, we obtained the ‘decoding accuracy’, where a more positive value suggests a higher pattern similarity between similar directions than between dissimilar directions (Wolff, Jochim, Akyürek, & Stokes, 2017). This process was repeated for 1000 times. The averaged results for each trial was used for the statistical test. The significance of decoding was assessed by comparing whether the decoding accuracy was different from zero. Cluster-based permutation was performed on the decoding accuracy time-series over participants (alpha = 0.05, cluster-based nonparametric alpha = 0.05, cluster statistic = sum, two-tail, permutation times = 10000, Maris & Oostenveld, 2007).

### Results

As shown in Fig. 1B, the speed-change detection performance was good when the target cue was presented (d’= 2.9+1.1, RT = 460+28ms). There was a trend of gaze shift towards the cued direction after the cue onset, during the delay and coherent moving period (see the gaze heatmap in Supplementary Fig. 1). To reveal the temporal characteristics of the feature-based gaze bias, we performed the Mahalanobis decoding on the gaze position data. Participants’ attention was biased toward the cued direction and this bias lasted during the entire delay period and further increased once the coherent moving started (Fig1.C, clustered *p* <.001, 0.164~2.4s). By contrast, little information about the distractor dots was obtained from the gaze data. These results were in agreement with the gaze heatmap which together suggested that gaze decoding can reveal the bias in attentional selection in the current feature-based task. Therefore, we examined the ocular dynamics under feature-based attentional inhibition with gaze decoding in Experiment 2.

## Experiment 2

### Participants

According to an estimation from G-power (Faul, Erdfelder, Lang, & Buchner, 2007), twenty-seven participants were supposed to be collected assuming a medium effect size (d = 0.5) and a high statistic power (80%) for a one-tail paired sample T-Test (alpha = 0.05). We tested twenty-seven participants in total and two participants were excluded as their performances were outliers to the sample (their d’ and RT were more than 2.5 SD beyond the mean). Twenty-five participants were included for the analysis (mean age = 21.7, range from 18 to 26, 8 males, all right-handed).

### Method

The general procedures were similar between Experiment 1 and Experiment 2. There were two conditions in Experiment 2. In the distractor-cueing condition, the cue indicated the moving direction of distractor dots. In the neutral condition, the cue was uninformative as neither the target dots nor the distractor dots would move towards the cued direction (see Fig. 2A). Since participants had no idea which group of dots would have the speed change in the neutral condition, they should attend both directions equally. Each condition had eight blocks of eighty trials and the blocks of the two conditions were randomly interleaved. Participants were informed of the cueing condition at the beginning of each block. There was a quiz at the end of the block asking participants to report which cueing condition they had performed. The overall accuracy was 97.5% suggesting that participants were aware of the cueing condition for each block.

**Figure 2.**
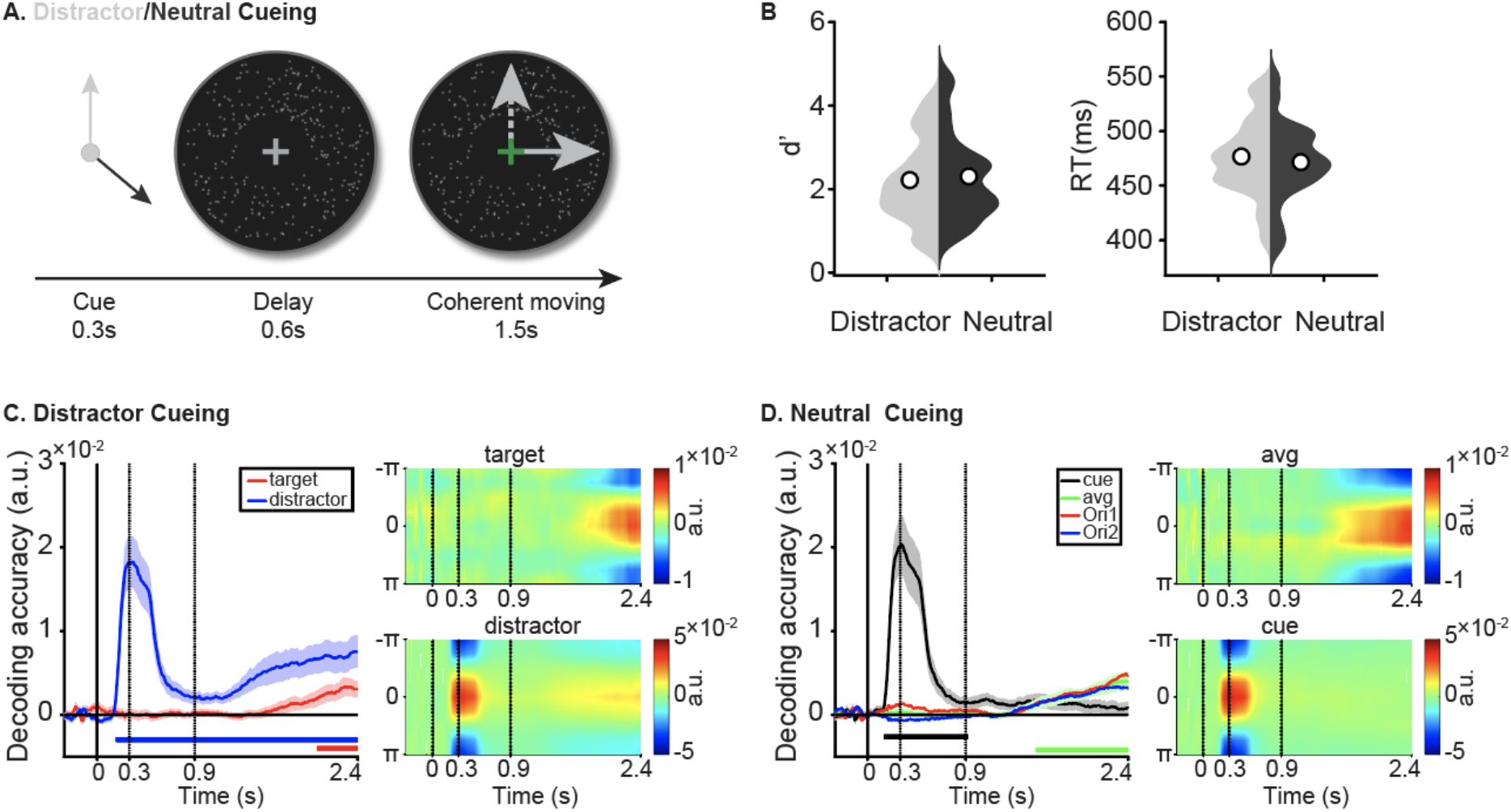
Task and results of Experiment 2. (A) Task. The cue in the distractor cueing condition (light arrow) indicated moving direction of the distractor dots, whereas neither the moving direction of the target dot nor the distractor dots will be informed by the cue (dark arrow) in the neutral condition (i.e., uninformative cue). (B) Behavioral performance of the two cueing conditions. Circles were the mean of all participants. (C) The direction decoding of distractor cueing condition. Solid lines below the x-axis are the cluster-permuted significant time intervals. (D) The direction decoding of neutral cueing condition. The green curve (avg) was the average of the red (ori1) and the blue curve (ori2).

### Data analysis

To examine whether distractor-cueing would benefit participants’ behavioral performance, the non-parametric permutation test (iteration = 100000) was performed on d’ and the averaged RTs of the two conditions. Direction decoding in the distractor cueing condition was similar to what we performed in Experiment 1. Because we had fewer trials for each direction bin, we partitioned the data into six-folds for the leave-one-out cross-validation. As for the decoding in neutral cueing condition, we arbitrarily labeled one direction as “ori1” and the other as “ori2” in each trial and performed decoding separately so that the decoding performance between the distractor and neutral condition would not be affected by the amount of trials. However, the statistical test was performed on the averaged decoding accuracy of the two directions.

### Results

There was no significant difference between distractor-cueing and neutral cueing conditions in the detection sensitivity (Mdis = 2.2+1.0, Mneu = 2.3+1.0, t(24) = −1.066, *p* =.759, Cohen’s d = .062) or RT (Mdis = 477+35ms, Mneu = 471+36ms, t(24) = 1.735, *p* = .611, Cohen’s d = .102). However, participants demonstrated distinct ocular dynamics under these two conditions. In the distractor-cueing condition, participants’ attention was biased by the distractor cue as informed by the decoding accuracy and the bias remained even after the coherent moving onset (Fig. 2C, blue curve, 0.17~2.4s, *p* < .001). In contrast, the target information gradually emerged until the late period of the coherent moving stage (Fig. 2C, red curve, 2.024~2.4s, *p* =.034). In the neutral cueing condition, although there was an initial bias cause by the uninformative cue (Fig. 2D, black curve, 0.15~0.926s, *p* =.002), participants disengaged from that direction and started to accumulate evidence for speed change at the two possible directions (Fig. 2D, green curve, 1.536~2.4s, *p* = .004).

## Experiment 3

Previous literature has suggested that distractor inhibition can be learned with extended practice. Suppression benefit would be found if the distractor feature remains the same throughout the experiment (Cunningham & Egeth, 2016; Moher, Lakshmanan, Egeth, & Ewen, 2014). In Experiment 3, we planned to investigate how the consistency of the distractor features affected the gaze dynamics.

### Participants

According to G-power (Faul et al., 2007), twenty-three participants were supposed to be collected to ensure a medium effect size (f = 0.25) and a high statistic power (85%) for a repeated one-factor ANOVA (alpha = 0.05). Twenty-three participants (mean age = 21.9, range from 18 to 26, 12 males, all right-handed) were recruited for the experiment.

### Method

Participants in Experiment 3 only completed the distractor cueing condition. However, instead of changing the distractor-cue every trial, we presented the same distractor cue across five consecutive trials. Target dots might change their moving directions in each trial. Participants performed 16 blocks and each block contained 80 trials.

### Data analysis

Repeated ANOVAs were performed on the d’ and RT to examine the behavioral difference across five repetitions. We first performed the decoding analysis on each repetition. The baseline correction was performed on each repetition separately to the mean of the pre-stimulus interval (−0.3~0s) of the repetition. Because we had fewer trials for each direction bin, the data was partitioned into five folds for the leave-one-out cross-validation. To reveal how expectations of the distractor direction modulated gazes, we performed the cross-repetition decoding. Five consecutive trials were baseline corrected to −0.3~0s of the first trial. Trials at each repetition were partitioned into five folds. We took the four folds of trials from each repetition to train a common classifier and then tested the remaining one-fold of trials from each repetition. This process was iterated for 1000 times.

To examine how repetition changed the decoding accuracy of target and distractor directions, we drew a best-fit line to account for the trend of mean decoding accuracies across five repetitions for each participant. Populations’ best-fit line slopes were tested against 0 using one-sample t-test (two-tailed, alpha = .05). A significant positive/negative t-value indicated that the decoding accuracy increased/decreased along with each repetition.

### Results

There were no significant differences among the five consecutive trials on d’ (F(4,88) =0.914, *p* = .460, η_p_^2^ = .040) or RT (F(4,88) = 0.861, *p* = .491, η_p_^2^ = .038). We speculated that the lack of the expected suppression benefits might be due to the relatively fewer repetitions in the current study compared with previous literature (e.g., Cunningham & Egeth, 2016; van Moorselaar & Slagter, 2019; Vatterott & Vecera, 2012; Wen et al., 2018).

Next, we investigated how repetitions of the distractor cue modulated the information about the target and distractor contained in the gaze positions. In terms of the averaged target direction decoding accuracy during the coherent moving stage, slopes of the best-fit lines were not statistically different from 0 (Fig.3B, t(22) = −0.15, *p* =.885, CI_95_ = [-.0011, .0009]). However, slopes of the best-fit lines for the decoding accuracy of distractor direction were marginally significant negative (Fig. 3C, t(22) = −2.01, *p* =.057, CI_95_ = [-.0031, 0]). Moreover, the averaged decoding accuracy between 0.3 to 2.4s of the fifth trial was smaller than the first trial (t(44) = 2.097, *p* = .021, one-tailed, alpha = .05).

**Figure 3.**
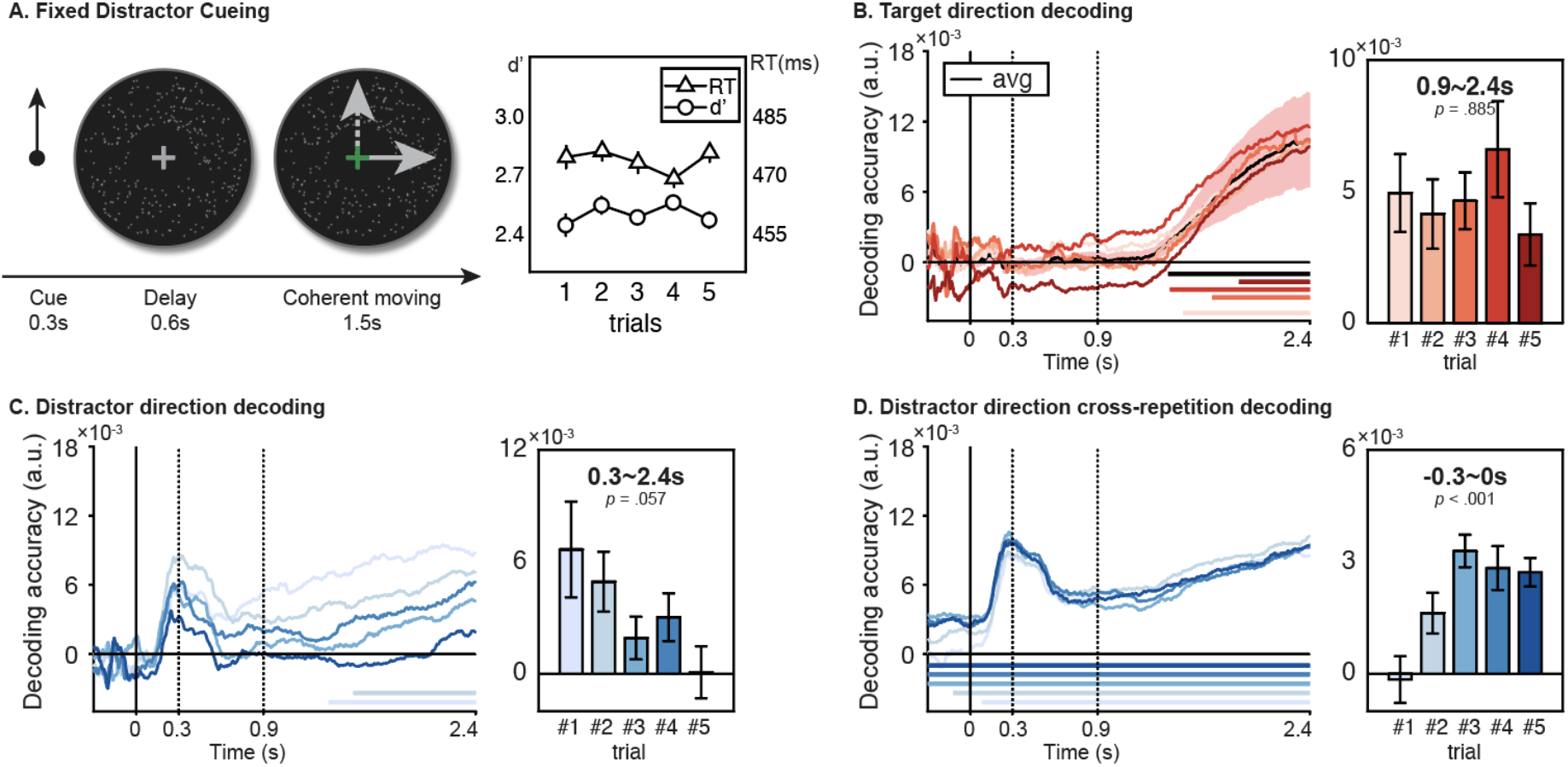
Task and results of Experiment 3. (A) Task and behavioral results. The cue indicated the moving direction of the distractor dots which remained unchanged in five consecutive trials. The dashed arrow in the coherent moving stage represented the moving direction of distractor dots while the solid arrow represented the target direction. The right panel showed the performance for speed-up detection. The errorbar represented the between-subject standard error. (B) Decoding of target direction for each trial repetition. The black curve is the averaged decoding accuracy across five repetitions. The solid lines below the x-axis indicate significant temporal clusters (trial 1, 1.506~2.4s, *p* = .007; trial 3, 1.712~2.4s, *p* = .014; trial 4, 1.412~2.4s, *p* = .009; trial 5, 1.906~2.4s, *p* = .035; average, 1.398~2.4s, *p* = .003). The right panel is the averaged decoding accuracy during the coherent moving stage (0.9~2.4s). Error bars represent within-subject standard errors. (C) Decoding of distractor direction for each trial repetition. Solid lines at the bottom represent the time intervals of significant clusters (trial 1, 1.344~2.4s, *p* = .012; trial 2, 1.52~2.4s, *p* = .024). The right panel shows the averaged decoding accuracy of distractor direction within the time window 0.3~2.4s. The *p*-value suggested that the population’s slopes of the best-fit line were marginally significant negative. (D) Cross-repetition decoding the distractor direction. (trial 1, 0.058~2.4s, *p* < .001; trial 2 −0.12~2.4s, *p* = .003; trial 3~5, −0.3~2.4s, *p* < .001). The right panel is the averaged decoding accuracy of the pre-stimulus interval (−0.3~0s). The *p*-value suggested that the population’s slopes of the best-fit line were significantly positive. See Supplementary Fig. S2-S4 for detailed decoding results at each repetition.

To demonstrated how the participants prepared the distractor inhibition under a predictive context, the cross-repetition decoding was conducted. The decoding accuracy of the pre-stimulus interval increased along with the repetition (Fig.3D, t(22) = 3.56, *p* < .001, CI_95_ = [.0003, .0011]). The reconstruction of the distractor direction in the fifth trial was larger than the first trial (t(44) = 3.009, *p* = .002, one-tailed, alpha = .05). This result suggested that, as the number of repetitions of the distractor cue increased, participants shifted their attention to the fixed distractor direction in advance before the start of the trials.

## Discussion

Distractor inhibition is challenging and context-dependent. Mixed results have been reported concerning whether distractors matched with the distractor cue could be efficiently suppressed. The current study investigated small gaze shifts in feature-based attentional control that could benefit from the inhibition of distractors in various cueing contingences and predictive contexts. Experiment 1 demonstrated that attention was biased toward the moving direction of the target dots and this bias could be quantitively measured through the decoding of gaze positions. In Experiment 2, participants were cued to the distractor’s moving direction and the results showed that participants’ attention was biased toward the cued distractor direction, as evidenced by decoded information from gaze positions. After being repeatedly cued to the same distractor direction in Experiment 3, participants strategically shifted their attention towards that direction proceeding the cue onset and gradually suppress further bias towards the distractor’s direction during the coherent moving stage.

In Experiment 2 where the distractor’s direction alternated on a trial-by-trial basis, participants’ attention was biased by the distractor cue and this bias lingered throughout the delay and coherent moving period. The sustained distractor information decoded from the gaze positions during the coherent moving stage suggested that distractor dots were still being attended and monitored. By contrast, cue-induced bias was quickly overcome in the neutral condition. Beck and colleagues (2018) examined the eye movement under the distractor cueing condition using a visual search task. They found that the first eye movement was biased towards the items that matched the to-be-ignored color cue but subsequent fixations were biased away from them. They further argued that configure an online feature-based negative template in a trial-by-trial manner is almost impossible and behavioral benefits of distractor cues emerge on condition that the later avoidance offsets the cost of early capture. Instead of measuring conventional ocular events (e.g., fixation and saccade), we performed direction decoding on the gaze positions to reveal the dynamics of feature-based distractor processing. Our results did not show the *later avoidance* as reported in Beck et al (2018) or the destroy component of the *search and destroy* process that was proposed by Moher and Egeth (2012). We conjectured that this discrepancy might be related to stimuli property and paradigm setting used in the present study. In a visual search task (Beck et al., 2018; Moher & Egeth, 2012), disengagement from the distractor location is the prerequisite to identify the target if the distractor captures attention. However, in the speed detection task used in the present study, the spatially intermingled moving dots of the target and distractor directions induced interference that was unlikely to be resolved by redirecting spatial attention. This interference occurred throughout the coherent moving stage, suggesting that sustained inhibition of the distractor direction could benefit the speed change detection in the target direction. Indeed, it has been suggested that distractors cannot be ignored before being selected in visual search (Donohue, Bartsch, Heinze, Schoenfeld, & Hopf, 2018). The decoding results in the distractor cueing condition demonstrated that monitoring the distractor direction while attending to the target direction serves as a means to reduce the between-direction interference.

In Experiment 3, when the distractor cue repeated in consecutive trials, the gaze dynamics manifested a functional dissociation at different stages. During the preparation, participants showed increased attention bias toward the distractor direction along with each repetition. Previous work showed that expectations could pre-activate the stimulus template (Blom, Feuerriegel, Johnson, Bode, & Hogendoorn, 2020; Kok, Mostert, & de Lange, 2017). Our results resonated with these findings and indicated that expectations could modulate gaze positions even before the trial started (see Fig. 3D). As a result of this strategical gaze-related attention shift, trial-wise re-deployment of attention to the distractor direction was not necessary as evidenced by the systematically deteriorated decoding performance during the coherent moving stage (see Fig. 3C). In van Moorselaar and Slagter (2019), they presented the distractor at the same place across trials to induce expectations about the upcoming distractor location. Their results suggested that expectations modulated distractor processing in a reactive manner as reflected by the smaller Pd component evoked by the distractor and lower decoding accuracy of the distractor location in the forth repeated trial relative to the initial trial. Indeed, we also found reduced reactive distractor inhibition as the repetition increased. Given the fixed distractor cue across consecutive five trials, our participants learned to shift their attention to the distractor’s direction proceeding the trial onset so as to avoid reactive processing of the distractor. Yet, they could still monitor the distractor with minimal attentional resources after stimulus onset. Taken together, the results of Experiment 3 suggested that predictive context facilitates distractor inhibition by strategically maintaining early attentional bias and reducing later switch costs for the monitoring process.

Instead of defining ocular events, we capitalized on the raw gaze positions to explore the real-time attentional bias in the present study. Because the participants were instructed to fixate at the central fixation cross and try to avoid tracking of the moving dots, the recorded data is treated as fixational eye movements. The fixational eye movement can be categorized as slow drift, microtremor, and microsaccades (Alexander & Martinez-Conde, 2019; Martinez-Conde, Macknik, & Hubel, 2004; Rucci & Poletti, 2015). The microsaccade has been regarded as an overt measure of covert attention while the slow drift is often assumed to random motions of the eye attempting to maintain visual fixations. However, one recent study suggested that the slow drift manifested stimulus-driven modulation in speed and direction (Malevich, Buonocore, & Hafed, 2020) and the changes in drift were behavior-relevant (Intoy & Rucci, 2020). We did not distinguish these two ocular events and performed direction decoding using the raw gaze data. This approach has been adopted by a recent study that showed that small gaze bias can reflect internal attention shifts in working memory (van Ede et al., 2019). Our results further demonstrate that the processes of feature-based attentional selection and inhibition could also be read-out from these small gaze shifts.

To conclude, we performed gaze decoding to reveal the dynamic process of feature-based attentional control. In the target cueing condition, tracking of the target direction was dominant and preserved to the end of the trial, whereas little information about the distractor could be read-out from the gaze positions. In the distractor cueing condition, distractor inhibition required constant monitoring of its direction, and this process was modulated by expectation. Expectations would promote distractor inhibition by biasing the gaze-related attention to the distractor’s direction in the preparatory stage and decreasing the cost of re-deployment of attention for distractor monitoring.

## Author Contributions

Wen Wen (Wen) developed the study concept and experimental design. Testing and data collection were performed by Wen and Yangming Zhang (Zhang). Wen performed the data analysis and interpretation under the supervision of Sheng Li (Li). Wen drafted the manuscript, and Li provided critical revisions. All authors approved the final version of the manuscript for submission.

## References

Alexander, R. G., & Martinez-Conde, S. (2019). Fixational Eye Movements. In C. Klein & U. Ettinger (Eds.), Eye Movement Research: An Introduction to its Scientific Foundations and Applications (pp. 73–115). https://doi.org/10.1007/978-3-030-20085-5_3

Arita, J. T., Carlisle, N. B., & Woodman, G. F. (2012). Templates for rejection: configuring attention to ignore task-irrelevant features. Journal of Experimental Psychology: Human Perception and Performance, 38(3), 580.

Awh, E., Belopolsky, A. V, & Theeuwes, J. (2012). Top-down versus bottom-up attentional control: A failed theoretical dichotomy. Trends in Cognitive Sciences, 16(8), 437–443.

Beck, V. M., & Hollingworth, A. (2015). Evidence for negative feature guidance in visual search is explained by spatial recoding. Journal of Experimental Psychology: Human Perception and Performance, 41(5), 1190–1196. https://doi.org/10.1037/xhp0000109

Beck, V. M., Luck, S. J., & Hollingworth, A. (2018). Whatever you do, don’t look at the. .: Evaluating guidance by an exclusionary attentional template. Journal of Experimental Psychology: Human Perception and Performance, 44(4), 645–662. https://doi.org/10.1037/xhp0000485

Becker, M. W., Hemsteger, S., & Peltier, C. (2015). No templates for rejection: A failure to configure attention to ignore task-irrelevant features. Visual Cognition, 23(9-10), 1150–1167. https://doi.org/10.1080/13506285.2016.1149532

Blom, T., Feuerriegel, D., Johnson, P., Bode, S., & Hogendoorn, H. (2020). Predictions drive neural representations of visual events ahead of incoming sensory information. Proceedings of the National Academy of Sciences of the United States of America, 117(13), 7510–7515. https://doi.org/10.1073/pnas.1917777117

Brainard, D. H. (1997). The psychophysics toolbox. Spatial Vision, 10(4), 433–436.

Conci, M., Deichsel, C., Müller, H. J., & Töllner, T. (2019). Feature guidance by negative attentional templates depends on search difficulty. Visual Cognition, 0(0), 1–10. https://doi.org/10.1080/13506285.2019.1581316

Cunningham, C. A., & Egeth, H. E. (2016). Taming the White Bear: Initial Costs and Eventual Benefits of Distractor Inhibition. Psychological Science, 27(4), 476–485. https://doi.org/10.1177/0956797615626564

Donohue, S. E., Bartsch, M. V., Heinze, H.-J., Schoenfeld, M. A., & Hopf, J.-M. (2018). Cortical Mechanisms of Prioritizing Selection for Rejection in Visual Search. The Journal of Neuroscience, 38(20), 4738–4748. https://doi.org/10.1523/jneurosci.2407-17.2018

Engbert, R., & Kliegl, R. (2003). Microsaccades uncover the orientation of covert attention. Vision Research, 43(9), 1035–1045.

Faul, F., Erdfelder, E., Lang, A.-G., & Buchner, A. (2007). G‪ Power 3: A flexible statistical power analysis program for the social, behavioral, and biomedical sciences. Behavior Research Methods, 39(2), 175–191.

Gaspar, J. M., & McDonald, J. J. (2014). Suppression of Salient Objects Prevents Distraction in Visual Search. Journal of Neuroscience, 34(16), 5658–5666. https://doi.org/10.1523/jneurosci.4161-13.2014

Gaspelin, N., Leonard, C. J., & Luck, S. J. (2015). Direct Evidence for Active Suppression of Salient-but-Irrelevant Sensory Inputs. Psychological Science, 26(11), 1740–1750. https://doi.org/10.1177/0956797615597913

Gaspelin, N., Leonard, C. J., & Luck, S. J. (2017). Suppression of overt attentional capture by salient-but-irrelevant color singletons. Attention, Perception, and Psychophysics, 79(1), 45–62. https://doi.org/10.3758/s13414-016-1209-1

Gaspelin, N., & Luck, S. J. (2018a). Distinguishing among potential mechanisms of singleton suppression. Journal of Experimental Psychology: Human Perception and Performance, 44(4), 626–644. https://doi.org/10.1037/xhp0000484

Gaspelin, N., & Luck, S. J. (2018b). The Role of Inhibition in Avoiding Distraction by Salient Stimuli. Trends in Cognitive Sciences, 22(1), 79–92. https://doi.org/10.1016/j.tics.2017.11.001

Gaspelin, N., & Luck, S. J. (2019). Inhibition as a potential resolution to the attentional capture debate. Current Opinion in Psychology, 29, 12–18. https://doi.org/10.1016/j.copsyc.2018.10.013

Geng, J. J. (2014). Attentional Mechanisms of Distractor Suppression. Current Directions in Psychological Science, 23(2), 147–153. https://doi.org/10.1177/0963721414525780

Geng, J. J., Won, B. Y., & Carlisle, N. B. (2019). Distractor Ignoring: Strategies, Learning, and Passive Filtering. Current Directions in Psychological Science. https://doi.org/10.1177/0963721419867099

Hafed, Z. M., & Clark, J. J. (2002). Microsaccades as an overt measure of covert attention shifts. Vision Research, 42(22), 2533–2545.

Han, S. W., & Kim, M. S. (2009). Do the Contents of Working Memory Capture Attention? Yes, But Cognitive Control Matters. Journal of Experimental Psychology: Human Perception and Performance, 35(5), 1292–1302. https://doi.org/10.1037/a0016452

Intoy, J., & Rucci, M. (2020). Finely tuned eye movements enhance visual acuity. Nature Communications, 11(1), 1–11.

Kok, P., Mostert, P., & de Lange, F. P. (2017). Prior expectations induce prestimulus sensory templates. Proceedings of the National Academy of Sciences, 114(39), 10473–10478. https://doi.org/10.1073/pnas.1705652114

Liesefeld, H. R., Liesefeld, A. M., Töllner, T., & Müller, H. J. (2017). Attentional capture in visual search: Capture and post-capture dynamics revealed by EEG. NeuroImage, 156(January), 166–173. https://doi.org/10.1016/j.neuroimage.2017.05.016

Macmillan, N. A., & Kaplan, H. L. (1985). Detection theory analysis of group data: estimating sensitivity from average hit and false-alarm rates. Psychological Bulletin, 98(1), 185.

Malevich, T., Buonocore, A., & Hafed, Z. M. (2020). Rapid stimulus-driven modulation of slow ocular position drifts. BioRxiv.

Maris, E., & Oostenveld, R. (2007). Nonparametric statistical testing of EEG-and MEG-data. Journal of Neuroscience Methods, 164(1), 177–190.

Martinez-Conde, S., Macknik, S. L., & Hubel, D. H. (2004). The role of fixational eye movements in visual perception. Nature Reviews Neuroscience, 5(3), 229–240.

Moher, J., & Egeth, H. E. (2012). The ignoring paradox: Cueing distractor features leads first to selection, then to inhibition of to-be-ignored items. Attention, Perception, and Psychophysics, 74(8), 1590–1605. https://doi.org/10.3758/s13414-012-0358-0

Moher, J., Lakshmanan, B. M., Egeth, H. E., & Ewen, J. B. (2014). Inhibition Drives Early Feature-Based Attention. Psychological Science, 25(2), 315–324. https://doi.org/10.1177/0956797613511257

Noonan, M. A. P., Crittenden, B. M., Jensen, O., & Stokes, M. G. (2018). Selective inhibition of distracting input. Behavioural Brain Research, 355(April), 36–47. https://doi.org/10.1016/j.bbr.2017.10.010

Olivers, C. N. L. (2009). What drives memory-driven attentional capture? The effects of memory type, display type, and search type. Journal of Experimental Psychology: Human Perception and Performance, 35(5), 1275.

Posner, M. I. (1980). Orienting of attention. Quarterly Journal of Experimental Psychology, 32(1), 3–25.

Rucci, M., & Poletti, M. (2015). Control and functions of fixational eye movements. Annual Review of Vision Science, 1, 499–518.

Sawaki, R., & Luck, S. J. (2011). Active suppression of distractors that match the contents of visual working memory. Visual Cognition, 19(7), 956–972. https://doi.org/10.1080/13506285.2011.603709

Soto, D., Heinke, D., Humphreys, G. W., & Blanco, M. J. (2005). Early, involuntary top-down guidance of attention from working memory. Journal of Experimental Psychology: Human Perception and Performance, 31(2), 248–261. https://doi.org/10.1037/0096-1523.31.2.248

Stilwell, B. T., & Vecera, S. P. (2019). Cued distractor rejection disrupts learned distractor rejection. Visual Cognition, 0(0), 1–16. https://doi.org/10.1080/13506285.2018.1564808

Tanda, T., & Kawahara, J. I. (2019). Association between cue lead time and template-for-rejection effect. Attention, Perception & Psychophysics.

Theeuwes, J. (2010). Top-down and bottom-up control of visual selection. Acta Psychologica, 135(2), 77–99. https://doi.org/10.1016/j.actpsy.2010.02.006

Töllner, T., Conci, M., & Müller, H. J. (2015). Predictive distractor context facilitates attentional selection of high, but not intermediate and low, salience targets. Human Brain Mapping, 36(3), 935–944.

van Ede, F., Chekroud, S. R., & Nobre, A. C. (2019). Human gaze tracks attentional focusing in memorized visual space. Nature Human Behaviour, 3(5), 462–470. https://doi.org/10.1038/s41562-019-0549-y

van Moorselaar, D., & Slagter, H. A. (2019). Learning What Is Irrelevant or Relevant: Expectations Facilitate Distractor Inhibition and Target Facilitation through Distinct Neural Mechanisms. The Journal of Neuroscience : The Official Journal of the Society for Neuroscience, 39(35), 6953–6967. https://doi.org/10.1523/JNEUROSCI.0593-19.2019

van Moorselaar, D., & Slagter, H. A. (2020). Inhibition in selective attention. Annals of the New York Academy of Sciences, 1464(1), 204.

Vatterott, D. B., & Vecera, S. P. (2012). Experience-dependent attentional tuning of distractor rejection. Psychonomic Bulletin and Review, 19(5), 871–878. https://doi.org/10.3758/s13423-012-0280-4

Vickery, T. J., King, L.-W., & Jiang, Y. (2005). Setting up the target template in visual search. Journal of Vision, 5(1), 8.

Wang, B., & Theeuwes, J. (2018). Statistical regularities modulate attentional capture. Journal of Experimental Psychology. Human Perception and Performance, 44(1), 13–17. https://doi.org/10.1037/xhp0000472

Wen, W., Hou, Y., & Li, S. (2018). Memory guidance in distractor suppression is governed by the availability of cognitive control. Attention, Perception, & Psychophysics, 80(5), 1157–1168.

Wolfe, J. M. (1994). Guided search 2.0 a revised model of visual search. Psychonomic Bulletin & Review, 1(2), 202–238.

Wolff, M. J., Jochim, J., Akyürek, E. G., & Stokes, M. G. (2017). Dynamic hidden states underlying working-memory-guided behavior. Nature Neuroscience, 20(6), 864–871. https://doi.org/10.1038/nn.4546

Woodman, G. F., Carlisle, N. B., & Reinhart, R. M. G. (2013). Where do we store the memory representations that guide attention? Journal of Vision, 13(3), 1–1. https://doi.org/10.1167/13.3.1

Woodman, Geoffrey F, & Luck, S. J. (2007). Do the contents of visual working memory automatically influence attentional selection during visual search? Journal of Experimental Psychology: Human Perception and Performance, 33(2), 363.

