## supplementary for "Gaze dynamics of feature-based distractor inhibition under the prior-knowledge and expectation"

Figure 1. Heatmap of gaze density at the cue (0~0.3s), delay (0.3~0.9s), and coherent moving phase (0.9~2.4s). The gaze positions were baselined to the pre-stimulus interval (-0.3~0s). Then, gaze positions within the selected time window from all trials were mapped to a 100*100 grid (covering a 4°×4° region). We obtained the two-dimensional histograms of gaze positions in the grid for each direction. To remove common bias irrespective of moving directions, we subtracted the averaged gaze density map of the eight directions. The yellow solid line indicates the moving direction of the target dots.

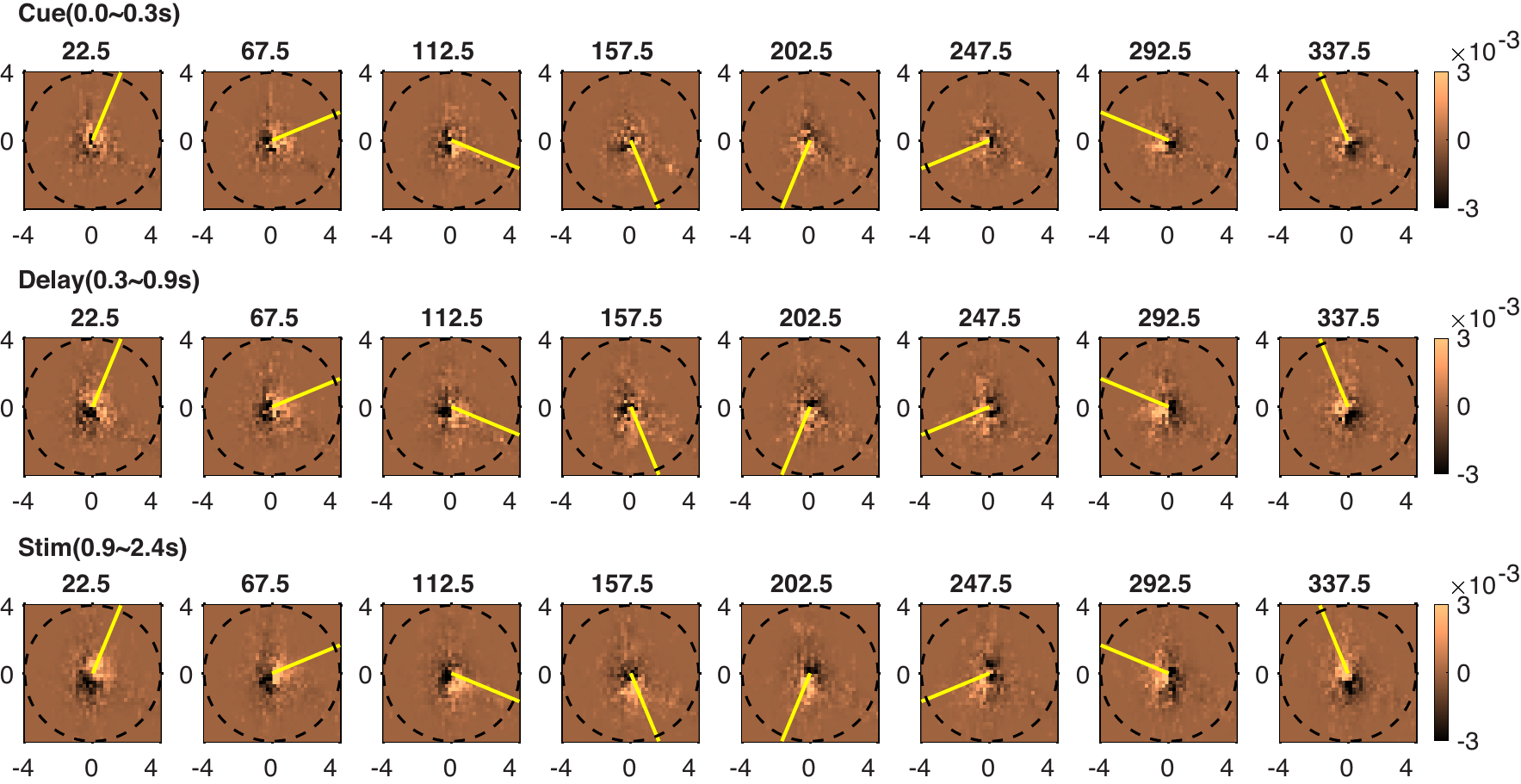

Figure 2. The direction decoding of target dots in Study 3. The shaded region represents SEM; Solid lines below the x-axis indicate significant clusters (*p* < 0.05).

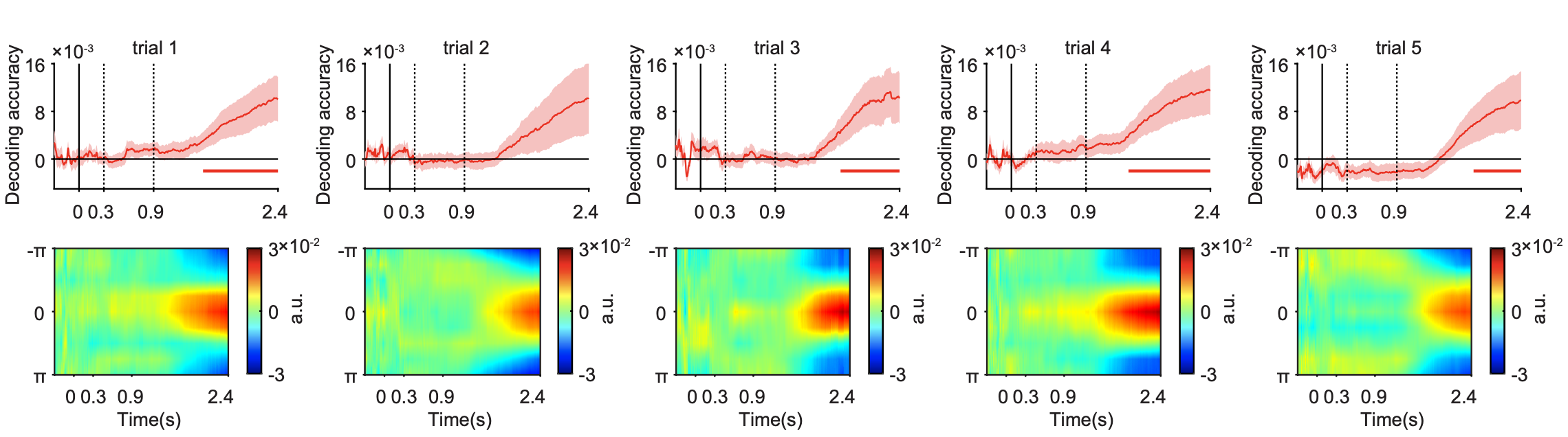

Figure 3. The direction decoding of distractor dots in Study 3. The shaded region represents SEM; Solid lines below the x-axis indicate significant clusters (*p* < 0.05).

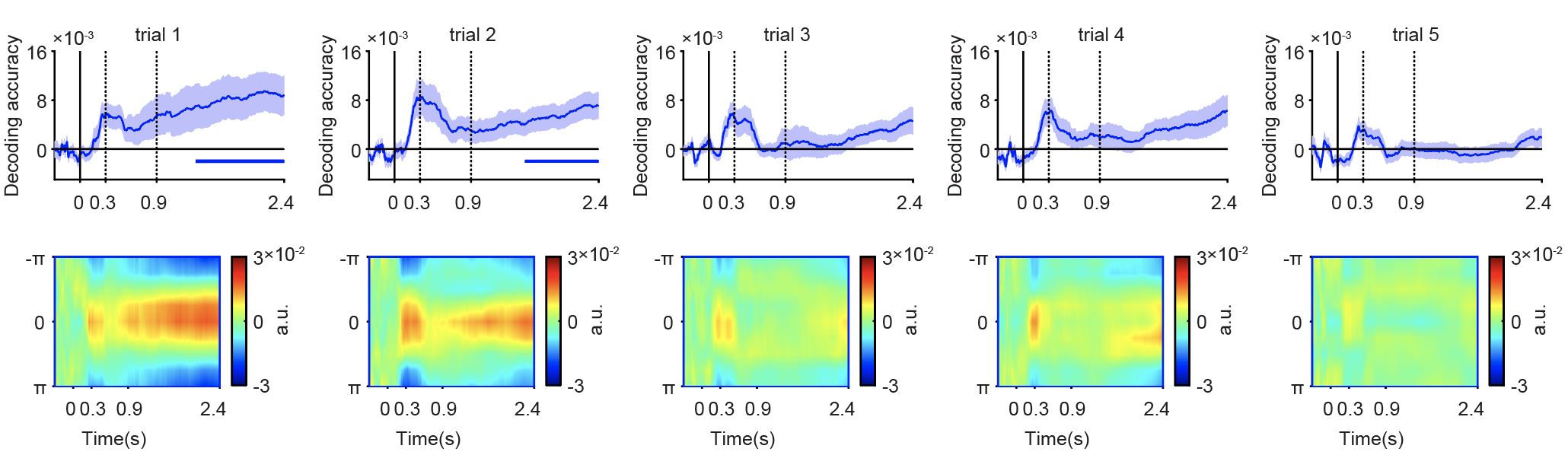

Figure 4. Cross-repetition decoding of moving direction of distractor dots in Study 3. The shaded region represents SEM; Solid lines below the x-axis indicate significant clusters (*p* < 0.05).

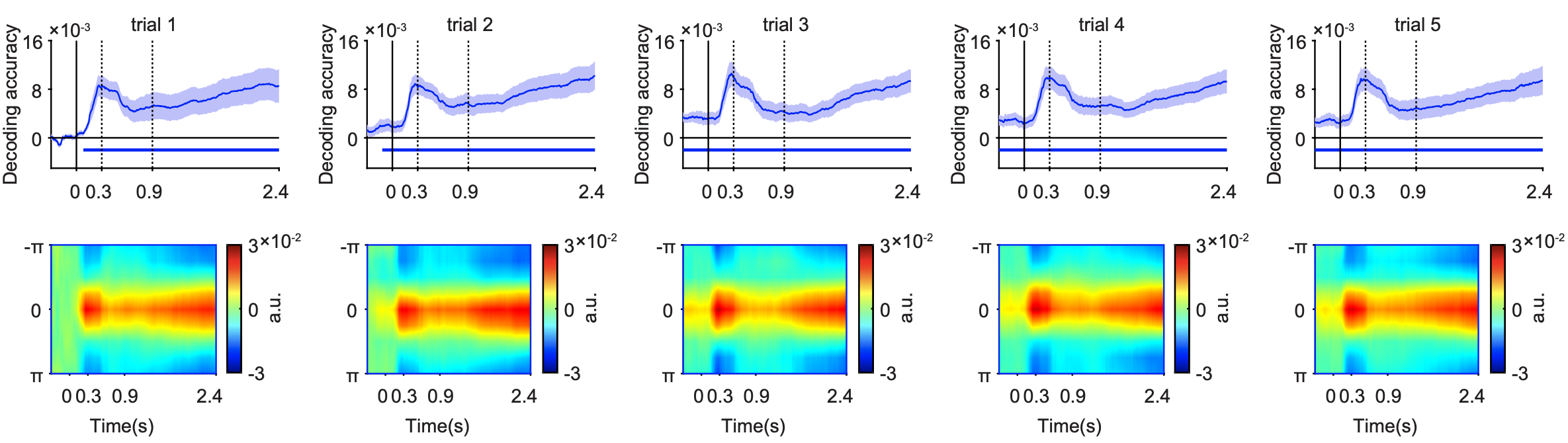
